# Reduced neutralization of SARS-CoV-2 B.1.617 variant by inactivated and RBD-subunit vaccine

**DOI:** 10.1101/2021.07.09.451732

**Authors:** Jie Hu, Xiao-yu Wei, Jin Xiang, Pai Peng, Feng-li Xu, Kang Wu, Fei-yang Luo, Ai-shun Jin, Liang Fang, Bei-zhong Liu, Kai Wang, Ni Tang, Ai-Long Huang

**Author notes:** These authors contributed equally to this work (Jie Hu, Xiao-yu Wei, Jin Xiang). **Corresponding authors:** Ai-long Huang, Ni Tang, Bei-zhong Liu, and Kai Wang, Key Laboratory of Molecular Biology for Infectious Diseases (Ministry of Education), Institute for Viral Hepatitis, Department of Infectious Diseases, The Second Affiliated Hospital, Chongqing Medical University, Chongqing, China.; (Ai-long Huang), (Ni Tang), (Bei-zhong Liu), (Kai Wang).

## Abstract

Coronavirus disease 2019 (COVID-19) is caused by severe acute respiratory syndrome coronavirus 2 (SARS-CoV-2). The Spike protein that mediates coronavirus entry into host cells is a major target for COVID-19 vaccines and antibody therapeutics. However, multiple variants of SARS-CoV-2 have emerged, which may potentially compromise vaccine effectiveness. Using a pseudovirus-based assay, we evaluated SARS-CoV-2 cell entry mediated by the viral Spike B.1.617 and B.1.1.7 variants. We also compared the neutralization ability of monoclonal antibodies from convalescent sera and neutralizing antibodies (NAbs) elicited by CoronaVac (inactivated vaccine) and ZF2001 (RBD-subunit vaccine) against B.1.617 and B.1.1.7 variants. Our results showed that, compared to D614G and B.1.1.7 variants, B.1.617 shows enhanced viral entry and membrane fusion, as well as more resistant to antibody neutralization. These findings have important implications for understanding viral infectivity and for immunization policy against SARS-CoV-2 variants.

## Introduction

The novel coronavirus reported in 2019 (2019-nCoV), officially named severe acute respiratory syndrome coronavirus 2 (SARS-CoV-2), is a new type of coronavirus belonging to the genus *Betacoronavirus*. It is a single-stranded RNA virus with a genome of approximately 29 Kb and with a high pathogenicity and high infectivity. As of July 8, 2021, there were more than 185 million confirmed cases of coronavirus disease 2019 (COVID-19) globally, including more than 4 million confirmed deaths (https://coronavirus.jhu.edu/). As SARS-CoV-2 continues to circulate in the human population, multiple mutations accumulate over time despite its proofreading capacity^1^. The Spike glycoprotein mutation D614G became dominant in SARS-CoV-2 during the early pandemic, which displayed increased infectivity and transmission^2^.

Spike-specific antibodies elicited by natural infection or vaccination contribute the majority of the neutralizing activity in human sera^3^. The receptor binding domain (RBD) in the S1 subunit of Spike protein binds to its cellular receptor angiotensin-converting enzyme 2 (ACE2) during viral entry, while the S2 subunit is required for the subsequent fusion of viral and cellular membranes^1^. Therefore, RBD is believed to be a major target of neutralizing antibodies (NAbs) and has been a focus of COVID-19 vaccine design^4,5^. Our previously studies showed that mutations in SARS-CoV-2 Spike protein could affect viral properties such as infectivity and neutralization resistance^6,7^. The newly emerged SARS-CoV-2 variant, B.1.617, first reported from India, which carries two mutations (L452R and E484Q) in its RBD is of particular concern. The AstraZeneca ChAdOx1 nCoV-19 vaccine appeared less effective than the Pfizer–BioNTech (BNT162b2) mRNA vaccine in preventing infection of SARS-CoV-2 B.1.617 variant^8^. Although mRNA-based COVID-19 vaccines provide above 90% efficacy against original SARS-CoV-2 strain, breakthrough infections with SARS-CoV-2 variants occur^9,10^. However, the efficacy of inactivated and RBD-subunit vaccines against B.1.617 variant is still unknown.

In this study, we used SARS-CoV-2 pseudovirus system to compare the viral entry efficiency *in vitro*, as well as the neutralization activities of convalescent sera, monoclonal antibodies (mAbs) and COVID-19 vaccine-elicited sera against these newly emerging SARS-CoV-2 variants, including the highly transmissible variants B.1.1.7, and B. 1.617.

## Materials and Methods

### Cell culture

HEK 293T (ATCC CRL-3216) and A549 cells (ATCC CCL-185) were purchased from the American Type Culture Collection (ATCC, Manassas, VA, USA). Cells were maintained in Dulbecco’s modified Eagle medium (DMEM; Hyclone, Waltham, MA, USA) supplemented with 10% fetal bovine serum (FBS; Gibco, Rockville, MD, USA), and 1% penicillin–streptomycin at 37 °C in 5% CO_2_. HEK 293T cells or A549 cells transfected with human ACE2 (293T-ACE2 or A549-ACE2) were cultured under the same conditions with the addition of G418 (0.5 mg/mL) to the medium.

### Sera samples

Convalescent sera samples from 20 patients with COVID-19 obtained in February and October 2020 at Yongchuan Hospital of Chongqing Medical University were previously reported.^11^ All sera were tested positive using magnetic chemiluminescence enzyme immunoassay (MCLIA) kits supplied by BioScience Co. (Tianjin, China)^12^. Patient sera were incubated at 56 °C for 30 min to inactivate the complement prior to experiments. Twenty CoronaVac vaccinee sera were obtained 7-14 days following the second dose of vaccine. Eight ZF2001 (RBD-subunit) vaccinee sera were obtained 26-30 days after booster immunization (second dose), and two ZF2001 vaccinee sera were obtained 14 days following the third dose of vaccine. The study was approved by the Ethics Commission of Chongqing Medical University (ref. no. 2020003). Written informed consent was waived by the Ethics Commission of the designated hospital for emerging infectious diseases.

### Plasmids and antibodies

The codon-optimized gene encoding reference strain (GenBank: QHD43416) SARS-CoV-2 Spike protein with C-terminal 19-amino acid deletion was synthesized by Sino Biological Inc (Beijing, China), and cloned into pCMV3 vector. D614G mutation was introduced using site-directed mutagenesis (denoted as pCMV3-S-D614G). SARS-CoV-2 B.1.617 and B.1.1.7 variant Spikes were codon-optimized and synthesized by GenScript Inc (Nanjing, China) and cloned into pCMV3 vector. The HIV-1 NL4-3 ΔEnv Vpr luciferase reporter vector (pNL4-3.Luc.R-E-) constructed by N. Landau^13^ was provided by Prof. Cheguo Cai from Wuhan University (Wuhan, China). The expression plasmid for human ACE2 was obtained from GeneCopoeia (Guangzhou, China). Anti-RBD monoclonal antibodies (mAbs) against the SARS-CoV-2 Spike protein were obtained from the blood samples of COVID-19 convalescent patients as described previously.^14^

### Production and titration of SARS-CoV-2 pseudoviruses

SARS-CoV-2 Spike pseudotyped viruses were produced as previously described with some modifications^15,16^. In brief, 5 × 10^6^ HEK 293T cells were co-transfected with pNL4-3.Luc.R-E- and recombinant SARS-CoV-2 Spike (D614G) plasmid or its derivatives (B.1.1.7 and B.1.617) using Lipofectamine 3000 (Invitrogen, Carlsbad, CA, USA). Supernatants containing pseudotyped viruses were harvested 48 h post-transfection, centrifuged, filtered through a 0.45-μm filter, and stored at −80°C. The titers of pseudoviruses were calculated by determining the number of viral RNA genomes per mL of viral stock solution using RT-qPCR with primers targeted the LTR^17^. Briefly, viral RNAs were extracted using TRIzol (Invitrogen, Rockville, MD, USA) and treated with RNase-free DNase (Promega, Madison, WI, USA) and re-purified using mini columns. Then, the RNA was amplified using the TaqMan One-Step RT-PCR Master Mix Reagents (Applied Biosystems, Thermo Fisher). A known quantity of pNL4-3.Luc.R-E-vector was used to generate standard curves. The prepared pseudovirues were adjusted to the same titer (copies/mL) for the following experiments.

### SARS-CoV-2 Spike-mediated pseudoviral entry assay

To detect Spike variant-mediated viral entry, 293T-ACE2 and A549-ACE2 cells (1.5× 10^4^) grown on 96-well plates were infected with 50 μL pseudoviruses (1 × 10^4^ copies). The cells were transferred to fresh DMEM medium 8 h post-infection, and RLU was measured 72 h post-infection using Luciferase Assay Reagent (Promega, Madison, WI, USA) according to the manufacturer’s protocol^18(p2)^.

### Cell-cell fusion assays

Syncytia formation assays were carried out as previously described with some modifications^19^. Briefly, plasmid pAdTrack-TO4-S, encoding SARS-CoV-2 Spike protein and enhanced green fluorescent protein (eGFP), was transfected into HEK 293T cells using Lipofectamine 3000 (Invitrogen). In parallel, another group of HEK 293T cells was transfected with hACE2 expressing plasmids. Two groups of cells were resuspended 24 h post-transfection, mixed a 1:1 ratio, and co-cultured in DMEM medium containing 10% FBS, 37 °C with 5% CO_2_, for 24 h, then observed the fusion under the fluorescence microscope.

### Western blot

To analyze Spike protein expression in cells, D614G, B.1.1.7, and B.1.617 variant Spike expressing plasmids were transfected into HEK 293T cells. Total protein was extracted from cells using radio immunoprecipitation assay Lysis Buffer (Beyotime, Shanghai, China) containing 1 mM phenylmethylsulfonyl fluoride (Beyotime). Equal amounts of protein samples were electrophoretically separated by 10% sodium dodecyl sulfate polyacrylamide gel electrophoresis, and then transferred to polyvinylidene difluoride membrane (Millipore, Billerica, MA, USA). The immunoblots were probed with the indicated antibodies. Protein bands were visualized using SuperSignal West Pico Chemiluminescent Substrate kits (Bio-Rad, Hercules, CA, USA) and quantified by densitometry using ImageJ software (NCBI, Bethesda, MD, USA).

### Pseudovirus-based neutralization assay

The 293T-ACE2 cells (1.5 × 10^4^ cells/well) were seeded on 96-well plates. For the neutralization assay, equivalent pseudoviruses (1 × 10^4^ copies in 50 μL) were incubated with serial dilutions of sera samples or mAbs for 1 h at 37 °C, then added to the 293T-ACE2 cells (with three replicates for each dilution). Luciferase activity was measured 72 h after infection. The titers of neutralizing antibodies were calculated as 50% inhibitory dose (ID_50_), the half-maximal inhibitory concentrations (IC_50_) of monoclonal antibodies (mAbs) against pseudoviruses was calculated using GraphPad Prism 8.0 software (GraphPad Software, San Diego, CA, USA).

### Statistical analyses

Statistical analyses of the data were performed using GraphPad Prism version 8.0 software. Quantitative data in histograms are shown as means ± SD. Statistical significance was determined using ANOVA for multiple comparisons. Student’s *t*-tests were applied to compare the two groups. Differences with *P* values < 0.05 were deemed statistically significant.

## Results

### B.1.617 variant Spike promotes viral entry and membrane fusion

Phylogenetic analysis showed that the newly emerged SARS-CoV-2 B.1.617 variant bearing common signature mutations G142D, L452R, E484Q, D614G and P681R, in its Spike glycoprotein (Fig. 1A). To assess the impact of these mutations on viral entry, synthetic codon-optimized B.1.617 and B.1.1.7 variant Spikes were cloned into mammalian expression vector respectively. Next, we generated pseudotyped SARS-CoV-2 using a lentiviral system, which introduced a Luc (luciferase) reporter gene for quantification of Spike-mediated viral entry. Thereafter, pNL4-3.Luc.R-E-was co-transfected with pS-D614G, pS-B.1.1.7 and pS-B.1.617 to package the Spike pseudotyped single-round Luc virus in HEK 293T cells. The titers of pseudoviruses were determined by reverse transcriptase quantitative polymerase chain reaction (RT-qPCR) expressed as the number of viral RNA genomes per mL, and then adjusted to the same concentration (1 × 10^4^ copies in 50 μL) for the following experiments.

**Figure 1.**
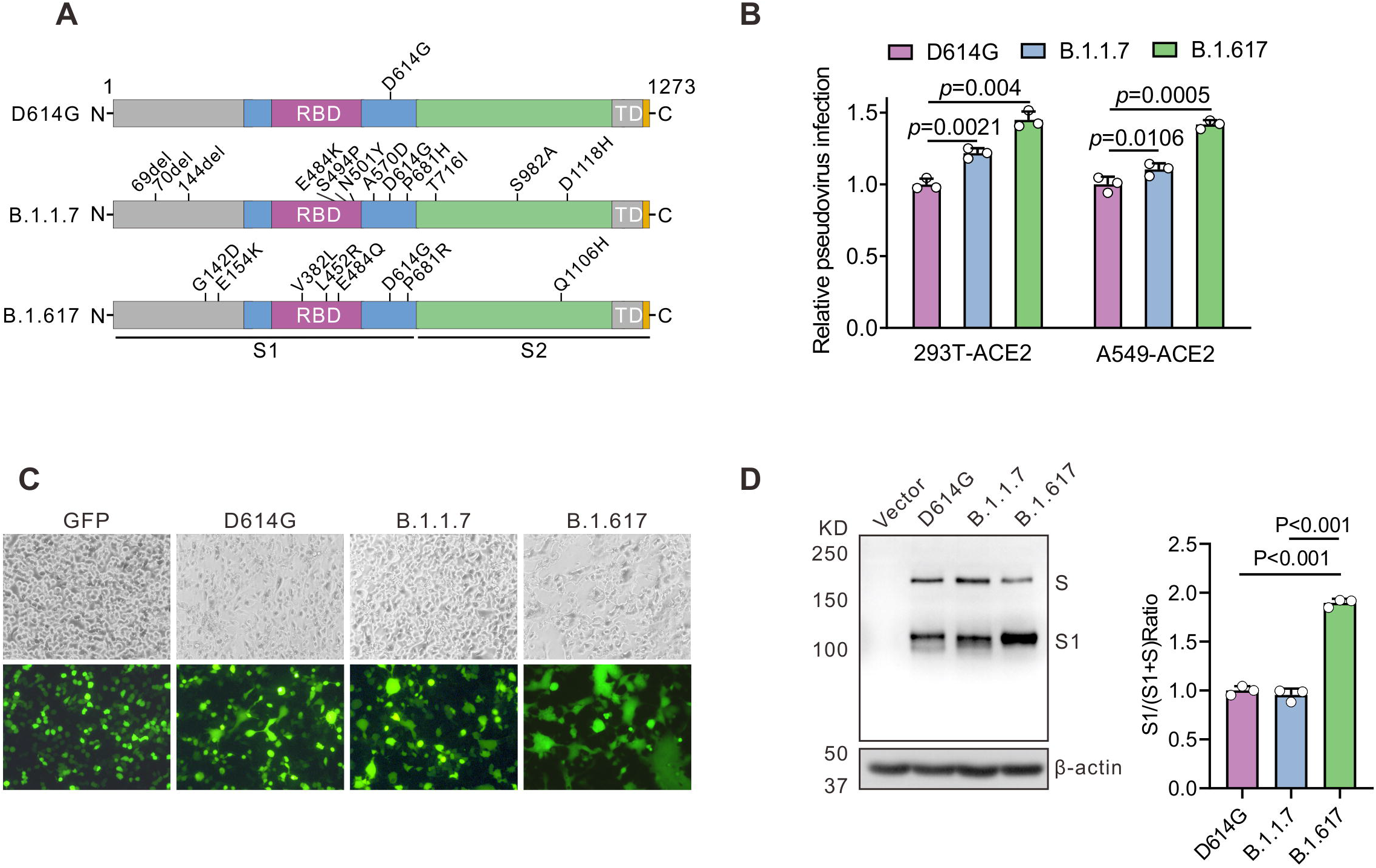
B.1.617 variant Spike protein of SARS-CoV-2 drives efficient viral entry and cell-cell fusion. (A) The diagram of SARS-CoV-2 Spike protein from D614G, B.1.1.7 and B.1.617 variants. D614G variant pseudovirus (containing the D614D mutation in Spike); B.1.1.7 variant pseudovirus (containing the H60/V70 and Y144 deletions and N501Y, A570D, D614G, P681H, T716I, S982A, and D1118H mutations in Spike); B.1.617 variant pseudovirus (containing the G142D, E154K, V382L, L452R, E484Q, D614G, P681R, and Q1106H mutations in Spike). (B) Infectivity of D614G, B.1.1.7 and B.1.617 variants pseudoviruses assessed in 293T-ACE2 and A549-ACE2 cells. Cells were inoculated with equivalent doses of each pseudotyped virus, at 6 h post inoculation, replaced the supernatants with fresh culture. Upon 72 h, cells were lysed with passive lysis buffer and analyzed the activity of firefly luciferase. (C) Quantitative cell-cell fusion assay. HEK293T cells expressing SARS-CoV-2 Spike variants D614G, B.1.1.7 and B.1.617 were mixed with ACE2-expressing target HEK293T cells (ratio 1: 1), and cell-cell fusion was analyzed by measuring the presence of syncytia by fluorescence microscopy. (D) Detection of Spike protein expression of D614G, B.1.1.7 and B.1.617 in HEK 293T cells by Western blot using the anti-RBD (receptor-binding domain) monoclonal antibody. To compare the S1 and S ratio, integrated density of S1/(S+S1) was quantitatively analyzed using ImageJ software. n = 3, ±SD. \*\**P* < 0.01.

The virus infectivity was determined by a Luc assay as previously described.^16^ As shown in Fig. 1B, to compare the viral entry efficiency meditated by Spike variants, we detected the Luc activity, the B.1.1.7 variant showed a slight increase in viral transduction over the D614G variant was 1.22-fold and 1.17-fold, while the B.1.617 variant over the D614G variant was 1.45-fold and 1.4-fold at 72 h post-infection in 293T-ACE2 and A549-ACE2 cells, respectively. These data suggest that the B.1.617 variant Spike protein significantly promotes viral entry into ACE2-expressing cells.

Next, we investigated Spike protein meditated cell-cell fusion. Coronavirus Spike protein on plasma membrane of effector cells can triggered its fusion of target cells (ACE2-expressing cells). B.1.617 variant Spike protein significantly increased fusion efficacy compared to D614G variant (Fig. 1C). To evaluate the expression and cleavage of SARS-CoV-2 Spike protein in a human cell line, the codon-optimized Spike-expressing plasmids (D614G, B.1.1.7 and B.1.617) were transfected into HEK 293T cells. The immunoblot analysis of whole cell lysates revealed that D614G, B.1.1.7 and B.1.617 Spike proteins showed two major protein bands (unprocessed S and cleaved S1 subunit), when allowed to react with the monoclonal antibody targeting the RBD on the SARS-CoV-2 Spike protein (Fig. 1D). However, the B.1.617-transfected cells showed a stronger S1 signal than D614G-transfected cells, indicating that the B.1.617 variant altered the cleavability of the Spike protein by cellular proteases. Collectedly, our data suggest that Spike protein of B.1.617 variant enhanced viral entry into ACE2-expressing cells and membrane fusion process, which may contribute to SARS-CoV-2 infectivity.

### Reduced neutralization by COVID-19 convalescent plasma

The plasma samples of 20 patients with COVID-19 obtained in February and October 2020 in Chongqing were previously reported.^11^ Using a luciferase-expressing lentiviral pseudotyping system, geometric mean titers (GMTs) were calculated to assess the neutralizing efficacy. The neutralizing activity of 5 samples against B.1.617 variant was reduced by >3-fold compared to D614G (Fig. 2A). Notably, the NAb titer of 6 samples (30%) was lower than the threshold against B.1.617 (Fig. 2A). 18 samples ID_50_ >40 against D614G pseudovirus, whereas the NAb titers of 3 samples (15%) and 6 samples (30%) decreased below the threshold against B.1.1.7 and B.1.617, respectively. The GMTs were 117 for D614G, 87 for B.1.1.7, and 50 for B.1.617 (Fig. 2B). These data indicate that B.1.1.7 and B.1.617 escape from neutralizing antibodies in some COVID-19 convalescent sera.

**Figure 2.**
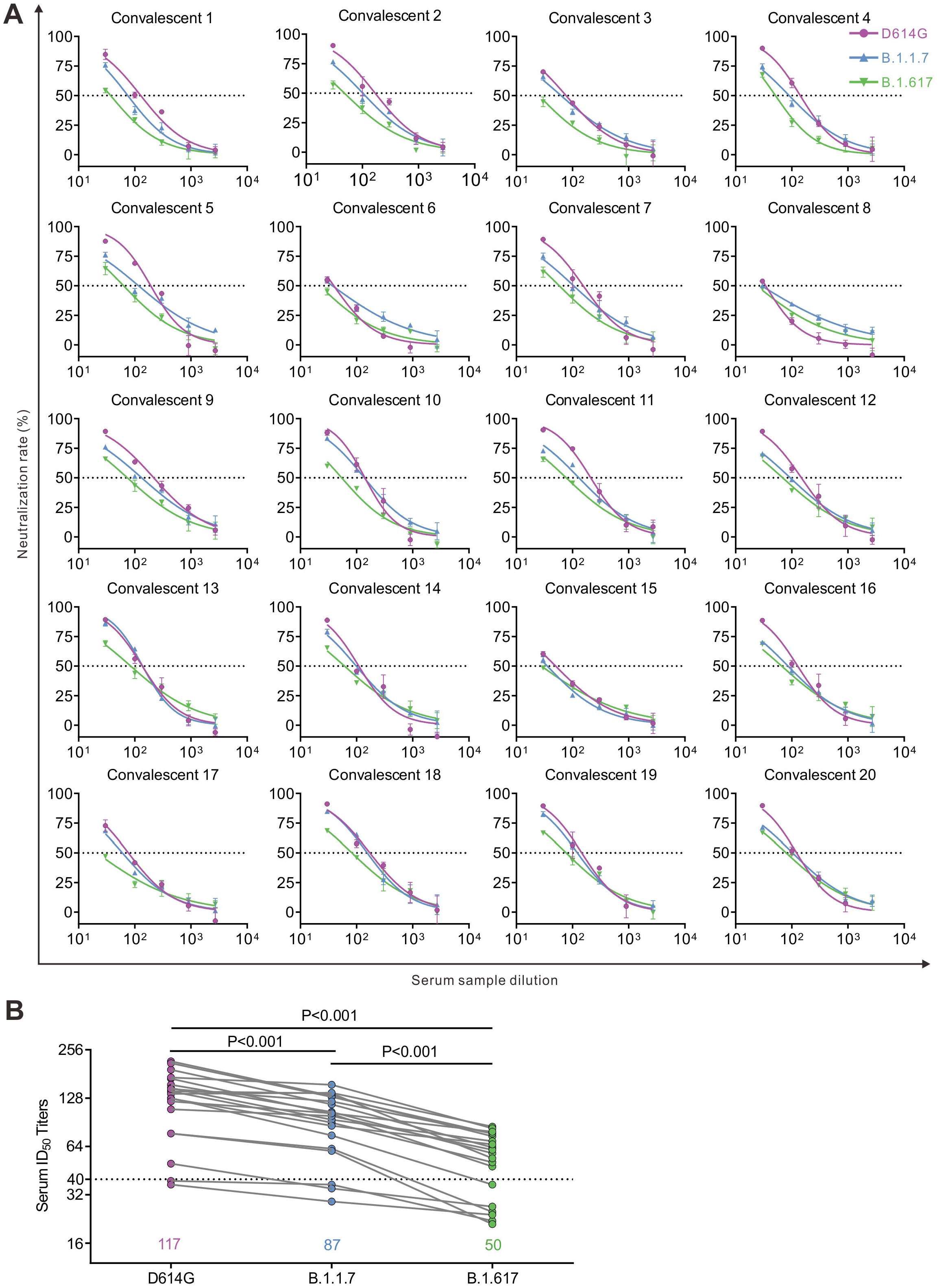
Neutralization efficiency of convalescent sera against D614G, B.1.1.7 and B.1.617 pseudotyped viruses. (A) Neutralizing activity of convalescent plasma (n=20) to D614G, B.1.1.7 and B.1.617 variants. Pseudotypes were incubated with different serum dilutions for 60 min at 37 °C, and then were added to the 293T-ACE2 cells. Upon 72 hours, cells were lysed with passive lysis buffer and analyzed the activity of firefly luciferase. The half-maximal neutralizing titer (ID_50_) was quantitatively analyzed using Graphpad 8.0.

### Resistance against monoclonal antibodies targeting the RBD

In addition, we assessed the impact of these variants on neutralizing activity of human monoclonal antibodies (mAbs) isolated from COVID-19 convalescent patients. Eight RBD-specific mAbs potent neutralizing SARS-CoV-2 obtained from the blood samples of COVID-19 convalescent patients were selected for this study.^14^ Among them, three mAbs showed less effective against B.1.1.7, and five against B.1.617 by 3-folds or more (Fig. 3A). Notably, the B.1.1.7 reduced the neutralization sensitivity with three mAbs (CQ012, CQ024 and CQ038) by 2 folds, and B.1.617 reduced the neutralization sensitivity with five mAbs (CQ012, CQ026, CQ038, CQ039 and CQ046) by 3 folds against D614G pseudovirus. Moreover, the B.1.617 reduced the neutralization sensitivity with the most potent mAb CQ046 by 4.6 folds, compared with that of D614G pseudovirus (Fig. 3B). The IC80 of mAb CQ046 decreased from 23.1 ng/ml (D614G) to 145.8 ng/ml (B.1.617). Together, both B.1.1.7 and B.1.617 reduced neutralization sensitivity to most mAbs tested. These data show the resistance of B.1.617 variant Spike proteins against monoclonal antibodies targeting the RBD.

**Figure 3.**
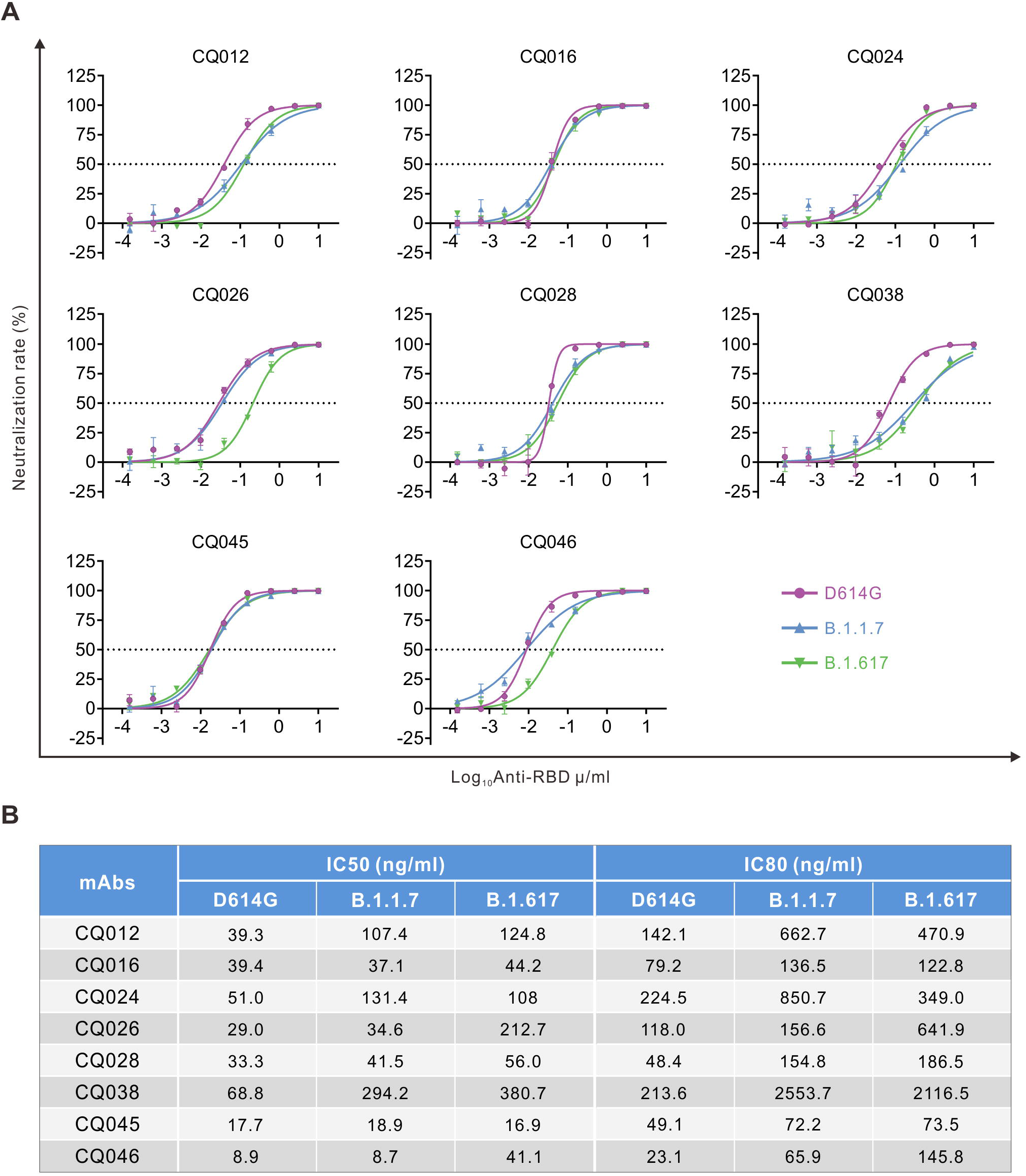
The RBD specific monoclonal antibodies (mAbs) against pseudoviruses. (A) The half-maximal inhibitory concentrations (IC_50_) representative neutralization curves for tested anti-RBD (receptor-binding domain) monoclonal antibodies (mAbs) against D614G, B.1.1.7 and B.1.617 pseudoviruses. Pseudotypes were incubated with different mAbs dilutions for 60 min at 37 °C, and then were incubated onto 293T-ACE2 cells. Upon 72 h, cells were lysed with passive lysis buffer and analyzed the activity of firefly luciferase. (B) The IC_50_ and IC_80_ were quantitatively analyzed using Graphpad 8.0.

### B.1.617 variant reduces sensitivity to vaccine-elicited antibodies

To evaluate the impact of the mutations present in Spike glycoprotein of SARS-CoV-2 variants on antibody neutralization, we compared the neutralization potency of COVID-19 vaccine-elicited antibodies against D614G, B.1.1.7 and B.1.617 Spike pseudotyed viruses. We collected serum from twenty individuals who received two doses of CoronaVac (inactivated vaccine) and eight individuals who received two doses of ZF2001 (RBD-subunit vaccine) vaccine, and two individuals who received three doses of ZF2001. Of the individuals who received three doses of ZF2001 (>14 days out from third dose) had robust neutralization of SARS-CoV-2 spike D614G, while those who received only two doses had lower but detectable neutralization (Fig. 4A-B). The GMT of ZF2001-elicited serum against the D614G, B.1.1.7 and B.1.617 were 151, 84, 49, respectively (Fig. 4C). Notably, ID_50_ of five samples against B.1.617 below the threshold were seen in two doses of ZF2001 sera. Together, B.1.617 showed more resistance to the neutralization of vaccinee serum than the D614G.These results indicate that it is of great importance to achieve three doses of ZF2001 vaccination.

**Figure 4.**
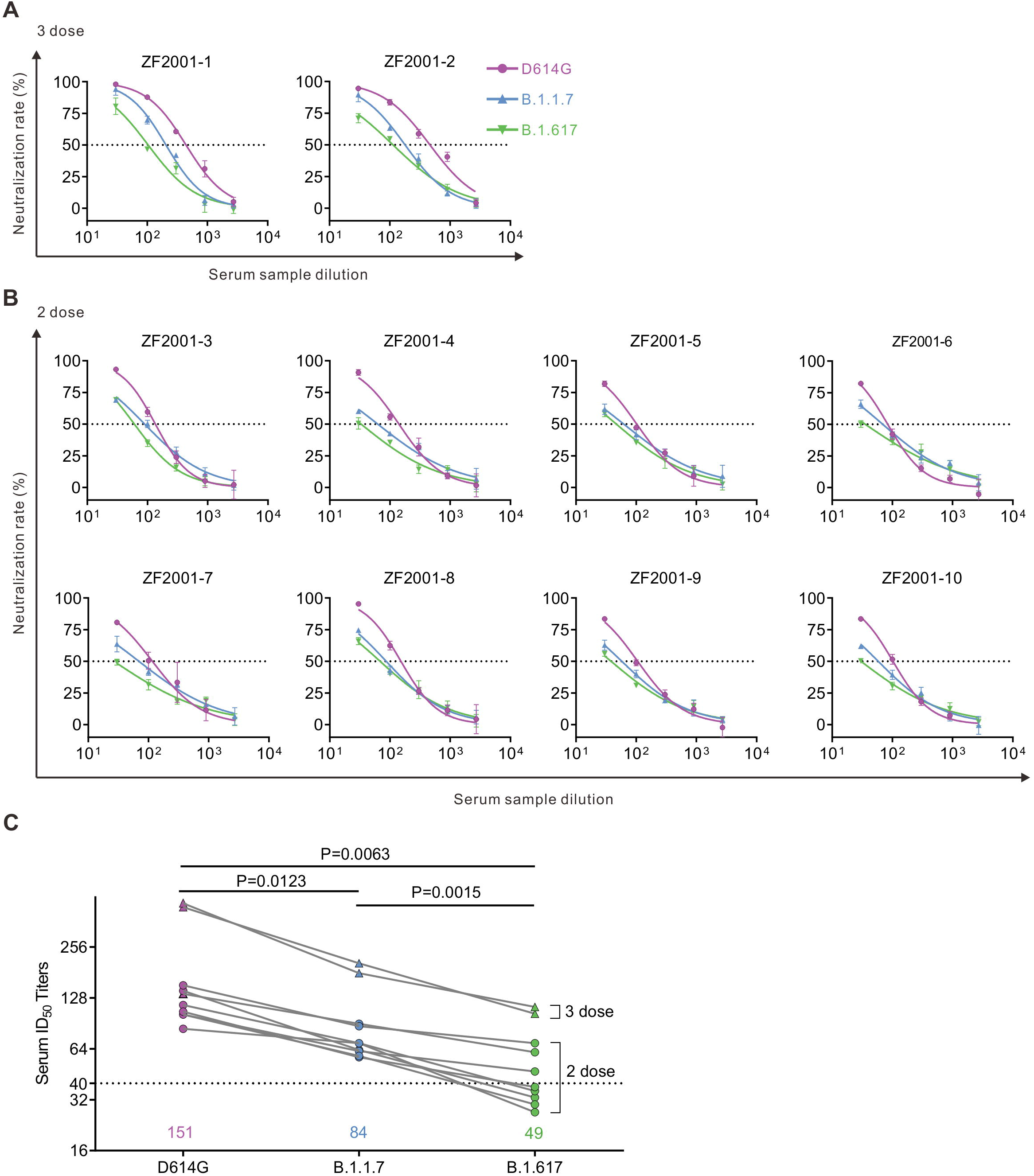
Detection of neutralizing antibodies against D614G, B.1.1.7 and B.1.617 pseudotyped viruses in ZF2001 vaccinee serum samples. (A-C) Neutralizing activity of ZF2001 (RBD-subunit vaccine) sera to D614G, B.1.1.7 and B.1.617. Two individuals who received three doses (A) and eight individuals who received two doses (B) of ZF2001. Pseudotypes were incubated with different serum dilutions for 60 min at 37 °C, and then were incubated onto 293T-ACE2 cells. Upon 72 h, cells were lysed with passive lysis buffer and analyzed the activity of firefly luciferase. The half-maximal neutralizing titer (ID_50_) was quantitatively analyzed using Graphpad 8.0. (C) The GMT of all ZF2001 vaccinee serum samples.

Nineteen CoronaVac-elicited vaccinees had substantial serum neutralizing activity against D614G Spike pseudotyped viruses (Fig. 5A). Compared with activity against the D614G, 35% (7/20) post-vaccination sera were decreased below the threshold against B.1.1.7, and 65% (13/20) were decreased below the threshold against B.1.617 (Fig. 5A). The average neutralization potency of the CoronaVac-elicited serum was reduced 2.5-fold for B.1.617 variant (GMT: 36) compared to D614G (GMT: 89) and reduced 1.6-fold for B.1.1.7 variant (GMT: 55) compared to D614G (GMT: 89) (Fig. 5B).

**Figure 5.**
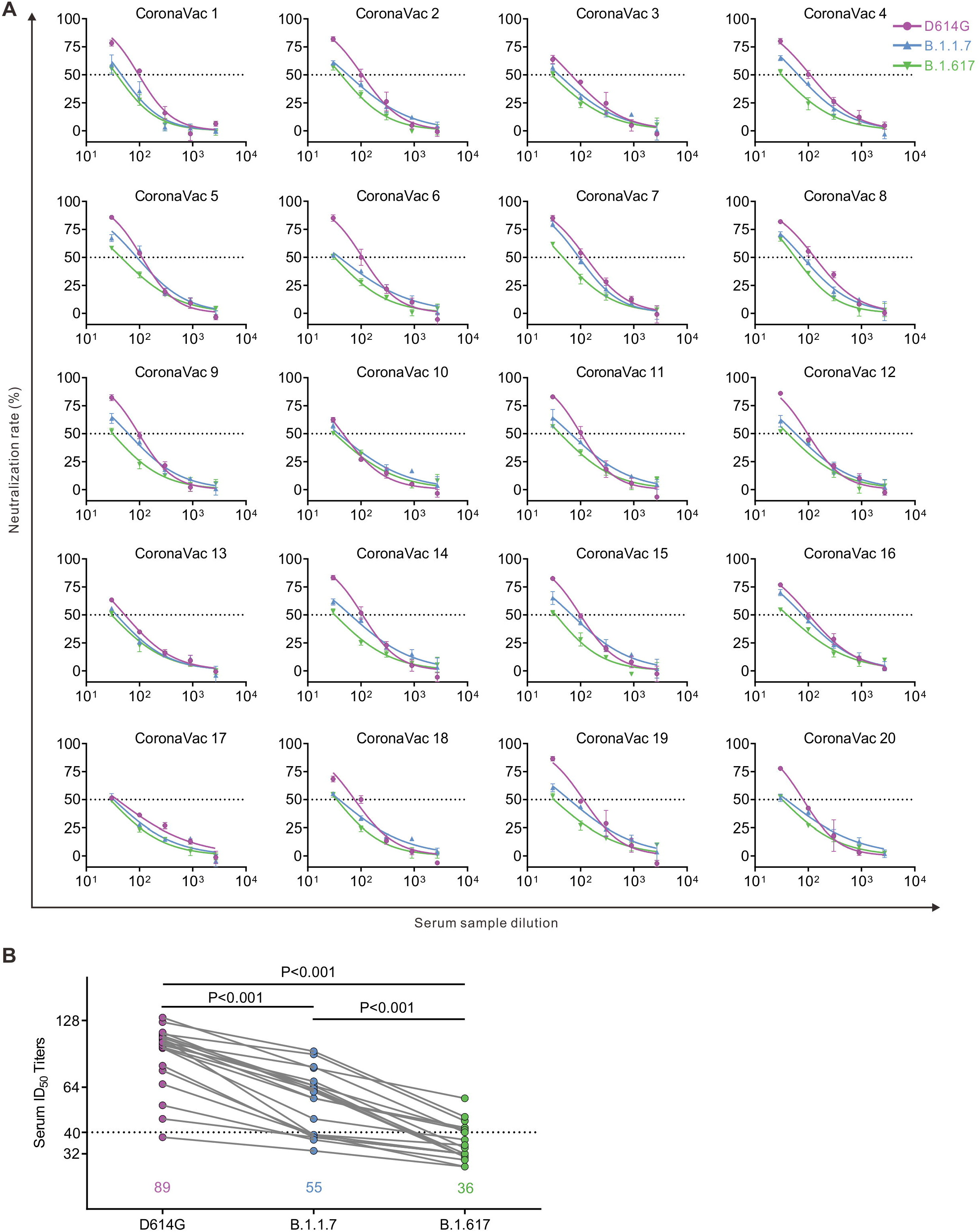
Detection of neutralizing antibodies against D614G, B.1.1.7 and B.1.617 pseudotyped viruses in CoronaVac vaccinee serum samples. (A) Neutralizing activity of CoronaVac (inactivated vaccine) sera (n=20) to D614G, B.1.1.7 and B.1.617. Pseudotypes were incubated with different serum dilutions for 60 min at 37 °C, and then were incubated onto 293T-ACE2 cells. Upon 72 h, cells were lysed with passive lysis buffer and analyzed the activity of firefly luciferase. The half-maximal neutralizing titer (ID_50_) was quantitatively analyzed using Graphpad 8.0. (B) The GMT of all CoronaVac vaccinee serum samples.

## Discussion

Due to the highly pathogenic nature of SARS-CoV-2, infectious SARS-CoV-2 must be handled in a biosafety level 3 (BSL-3) facility. Here, using luciferase-expressing lentiviral pseudotype system, we compared viral entry meditated by three SARS-CoV-2 Spike variants: the original D614G variant (identified during the first wave), B.1.1.7 variant (first detected in United Kingdom during the second wave), and B.1.617 variant first reported in India. Our data indicated that B.1.617 variant Spike promotes virus infectivity through enhanced viral entry and membrane fusion, which may play an important role in increased transmissibility of this variant. These findings are highly consistent with previous studies^20^. L452R mutation in the RBD was reported to increase SARS-CoV-2 infectivity and fusogenicity^21^. P681R, a highly conserved mutation in the B.1.617 lineages, also enhanced SARS-CoV-2 Spike-mediated cell-cell fusion^22^. At the time of preparing this manuscript, the B. 1.617.2 variant has displaced B.1.1.7 variant as the dominant SARS-CoV-2 strain in UK and other countries^23,24^.

Another explanation for the increased transmission of B. 1.617 variant might be the enhanced ability for the virus to evade immune system. In this study, we compared NAb titres of sera collected from previously SARS-CoV-2 infected individuals, CoronaVac (inactivated vaccine) and ZF2001 (RBD-subunit vaccine) vaccinated persons against three SARS-CoV-2 Spike variants. We found that B.1.617 variant Spike showed more resistant to antibody neutralization. B.1.617 reduced the neutralization of CoronaVac vaccine by 2.5 times, and ZF2001 vaccine by 3.1 times. Consistently, Liu et al reported that B.1.617 reduced the neutralization of convalescent plasma by 3.9 times, Pfizer-BioNTech vaccine by 2.7 times, and Oxford-AstraZeneca vaccine by 2.6 times^25^.

The RBD of the B.1.617 Spike contains two mutations, L452R and E484Q, which were thought to confer to immune evasion. Several studies have demonstrated that the E484K mutation in the RBD significantly reduced susceptibility to neutralization, as seen in B.1.351 (South Africa) and P.1 (Brazil) variants.^26–29^ E484Q mutation occurring in the same position as E484K, was also demonstrated to be associated with immune escape^25,30^. Another key mutation in the RBD of B.1.617 is L452R. Recent studies suggested that the L452R mutation of B.1.427/B.1.429 variant Spike also contributes to its escape from NAbs^31,32^.

The limitation of this study include its small sample size, only focus on pseudovirus-basded antibody neutralization in cell culture, and the possibility that mutations may alter neutralization by modulating Spike function rather than its antigenicity. To fully characterize the features of B.1.617 variant, *in vivo* study with authentic virus and the role of memory T or B cells in protection against this variant will be required. Conclusions about vaccine-mediated protection must be validated by real-world data collected in regions where B.1.617 variant is circulating.

Collectively, this study will be helpful for understanding the increased spread of B.1.617 variant and highlight the need to in depth survey of this variant. Given the evolving nature of the SARS-CoV-2 RNA genome, new variant of concern will continue to arise, which may threaten vaccine efficacy. Therefore, antibody therapeutics and vaccine evaluations against new variants are worthy of further investigation.

## Acknowledgements

We would like to thank Professor Cheguo Cai (Wuhan University, Wuhan, China) for providing the pNL4-3.Luc.R-E-plasmid. We also thank all the volunteers who participated in this research. This work was supported by the Natural Science Foundation Project of Chongqing (cstc2019jscx-dxwtBX0019), the Emergency Project from the Science & Technology Commission of Chongqing (cstc2020jscx-fyzx0053, cstc2020jscx-dxwtB0050), Kuanren Talents Program of the second affiliated hospital of Chongqing Medical University, the Emergency Project for Novel Coronavirus Pneumonia from the Chongqing Medical University (CQMUNCP0302), China Postdoctoral Science Foundation (2021M693924), and Chongqing Postdoctoral Science Special Foundation (2010010005216630).

